# Reactive Centre Loop Dynamics and Serpin Specificity

**DOI:** 10.1101/391482

**Authors:** Emilia M. Marijanovic, James Fodor, Blake T. Riley, Benjamin T. Porebski, Mauricio G. S. Costa, Itamar Kass, David E. Hoke, Sheena McGowan, Ashley M. Buckle

**Author notes:** Correspondence to: Ashley M. Buckle, Department of Biochemistry and Molecular Biology, Biomedicine Discovery Institute, Monash University, Victoria 3800, Australia.

## Abstract

Serine proteinase inhibitors (serpins), typically fold to a metastable native state and undergo a major conformational change in order to inhibit target proteases. However, conformational labiality of the native serpin fold renders them susceptible to misfolding and aggregation, and underlies misfolding diseases such as a1-antitrypsin deficiency. Serpin specificity towards its protease target is dictated by its flexible and solvent exposed reactive centre loop (RCL), which forms the initial interaction with the target protease during inhibition. Previous studies have attempted to alter the specificity by mutating the RCL to that of a target serpin, but the rules governing specificity are not understood well enough yet to enable specificity to be engineered at will. In this paper, we use *conserpin*, a synthetic, thermostable serpin, as a model protein with which to investigate the determinants of serpin specificity by engineering its RCL. Replacing the RCL sequence with that from α1-antitrypsin fails to restore specificity against trypsin or human neutrophil elastase. Structural determination of the RCL-engineered conserpin and molecular dynamics simulations indicate that, although the RCL sequence may partially dictate specificity, local electrostatics and RCL dynamics may dictate the rate of insertion during protease inhibition, and thus whether it behaves as an inhibitor or a substrate. Engineering serpin specificity is therefore substantially more complex than solely manipulating the RCL sequence, and will require a more thorough understanding of how conformational dynamics achieves the delicate balance between stability, folding and function required by the exquisite serpin mechanism of action.

## Introduction

Over 1,500 serpins have been identified to date. Inhibitory family members typically fold to a metastable native state that undergoes a major conformational change (termed the stressed [S] to relaxed [R] transition) central for the protease inhibitory mechanism^1^. The S to R transition is accompanied by a major increase in stability. The archetypal serpin fold is exemplified by α1-antitrypsin (α1-AT), a single domain protein consisting of 394 residues, which folds into 3 β-sheets (A C) and 9 α-helices (A I) that surround the central β-sheet scaffold^2^. The reactive center loop (RCL) protrudes from the main body of the molecule and contains the scissile bond (P1 and P1’ residues), which mediates α1 -AT’s inhibitory specificity against the target protease, neutrophil elastase (HNE).

The inhibitory mechanism of serpins is structurally well understood^1^. Briefly, a target protease initially interacts with and cleaves the RCL of the serpin. However, following RCL cleavage, but prior to the final hydrolysis of the acyl enzyme intermediate, the RCL inserts into the middle of the serpin’s β-sheet A to form an extra strand^1,3^. Since the protease is still covalently linked to the P1 residue, the process of RCL insertion results in the translocation of the protease to the opposite end of the molecule. In the final complex, the protease active site is distorted and trapped as the acyl enzyme intermediate^1,4^.

In certain circumstances the serpin RCL can spontaneously insert, either partially (delta conformation), or fully (latent conformation) into the body of the serpin molecule without being cleaved^5^. Both latent and delta conformations are considerably more thermodynamically stable than the active, native state although they are inactive as protease inhibitors. Folding to the latent conformation is thought to occur via a late, irreversible folding step that is accessible from the native or a highly native-like state^6,7^. As such, transition to the latent state can be triggered by perturbations to the native state via small changes in solution conditions such as temperature or pH^4,8,9^, or by spontaneous formation over long time scales^10,11^.

Human α1-AT is an extremely potent inhibitor of its target protease HNE, with a rate of association (kass) 6 × 10^7^ M^-1^ s^-1^, forming a serpin-protease complex that is stable for several days^12,13^. The metastable nature of α1-AT is required to facilitate the large conformational change required for its inhibitory function, and the rate of RCL insertion into β-sheet A is the main determinant of whether the acyl linkage between serpin and protease is maintained or disrupted. If RCL insertion is rapid, the inhibitory pathway proceeds. If the RCL insertion is too slow, the serpin becomes a substrate; the de-acylation step of the protease’s catalytic mechanism is complete and cleavage of the P1–P1’ bond occurs without protease inhibition. The cleaved, de-acylated RCL still inserts into the body of the serpin, resulting in an inactive inhibitor^14^.

Two regions of the RCL appear to govern inhibitory function and specificity. The first, a highly-conserved hinge region (resides P15–P9) consisting of short chain amino acids, facilitates RCL insertion into the A β-sheet. Mutations in the hinge region result in the serpin becoming a substrate rather than an inhibitor^15^. The second region is the P1 residue, thought to determine specificity towards a protease. Serpins with a P1 arginine (e.g. antithrombin III) are known to target proteases of the coagulation cascade, including thrombin and Factor Xa^16,17^. In α1-AT, mutation of P1 methionine to arginine (the Pittsburgh mutation), changes the specificity from HNE to thrombin, resulting in a bleeding disorder^18^.

Given the importance of the RCL, it has been the focus of previous attempts aimed at altering serpin specificity, via mutation of RCL residues or swapping RCL sequences between serpins. Chimeric serpins have been made between plasminogen activator inhibitor-1 (PAI-1) and antithrombin-III (ATIII)^19,20^, α1-AT and antithrombin-III^21,22^, α1-AT and ovalbumin^23^, and alpha1-antichymotrypsin (ACT) and α1-AT^13,24,25^. In all cases, specificity could only be transferred partially, as each chimera has a reduced second-order rate constant and a higher SI to a target protease in comparison to the original serpin. The most effective chimera produced, without a cofactor, was ACT with P3-P3’ of α1-AT. This chimera achieved an SI of 1.4 and a second-order rate constant (k’/[I]) of 1.1 10^5^ M^-1^ s’^1^, two orders of magnitude slower than that of α1-AT^13^. Therefore, it is highly likely that the determinants of specificity are more complex than the RCL region alone, and other regions may play a role, for example exosite interactions in the serpin–protease complex^20,26–28^

In previous work, we designed and characterized conserpin, a synthetic serpin that folds reversibly, is functional, thermostable and resistant to polymerization^29^. Conserpin was designed using consensus engineering, using a sequence alignment of 212 serpin sequences and determining the most frequently occurring amino acid residue at each position. Since it is thermostable and easier to produce in recombinant form, it is ideally suited as a model in protein engineering studies. Conserpin shares 59% sequence identity to α1-AT, with 154 residue differences scattered throughout the structure. Its RCL sequence is sufficiently different from all other serpins such that it no longer resembles an RCL of any serpin with a known target protease. A recent study that investigated the folding pathway of conserpin engineered the P7-P2’ sequence of α1-AT into its RCL^30^. The resulting conserpin/α1-AT chimera inhibits chymotrypsin with an SI of 1.46, however, no SI was calculated against HNE. The chimera forms a weak complex with HNE that is detectable using SDS-PAGE, however, the majority of the serpin molecules are cleaved without complex formation.

In this study, we have exploited the unique folding characteristics of conserpin and employ it as a model serpin with which to investigate the determinants of specificity. We investigated the effect of replacing the RCL of conserpin with the corresponding sequence from α1-AT on inhibitory specificity towards HNE. Here, the chimera molecule, called conserpin-AAT_RCL_, remains thermostable, yet despite possessing the RCL sequence of α1-AT, specificity against HNE was not restored to the extent of α1-AT. Structural analysis and molecular dynamics simulations indicate that specificity is also governed by other, complex factors involving RCL dynamics, and surface electrostatics of regions external to the RCL.

## Results

### Biophysical and functional characterisation of a conserpin/α1-AT chimera

With the aim of changing the specificity of conserpin to that of α1-AT, a conserpin/α1-AT chimera was previously produced^30^, where 9 residues within the RCL (P7-P2’) were swapped with the corresponding residues from α1-AT (Fig. 1A). The resulting chimera, conserpin-AAT_RCL_ (379 aa) has a 61% sequence identity with α1-AT (148 residue differences). Conserpin-AAT_RCL_ was expressed in *E. coli* and purified from the soluble fraction by affinity and size exclusion chromatography as described previously^29^. We first investigated the biophysical properties of conserpin-AAT_RCL_ to ensure that swapping the RCL did not alter them. The majority of serpins irreversibly unfold upon heating with a midpoint temperature transition (T_m_) of ~55-65°C^31–33^. Using variable temperature far-UV circular dichroism (CD) to measure the thermostability, conserpin-AAT_RCL_ was heated from 35 to 95°C at a rate of 1 °C /min, and upon reaching 95°C, an incomplete transition to the unfolded state was observed. Following a subsequent 1 °C /min decrease in temperature from 95 to 35°C, minute changes in signal was observed (Fig. 1B). In addition, far-UV spectral scans before and after thermal unfolding showed minute differences in the signals, suggesting the absence of a large heat-induced conformational change (Fig. 1C). Complete unfolding was only achieved in the presence in 2 M guanidine hydrochloride (GdnHCl) with a T_m_ of 72.2 ± 0.1°C. Visible precipitation was observed upon cooling from 95 to 35°C was not observed (Fig. 1D). Thus, high thermostability is consistent with that of parent conserpin molecule^29^ and indicates that incorporation of the α1-AT RCL does not reduce the thermostability of the conserpin scaffold.

**Figure 1.**
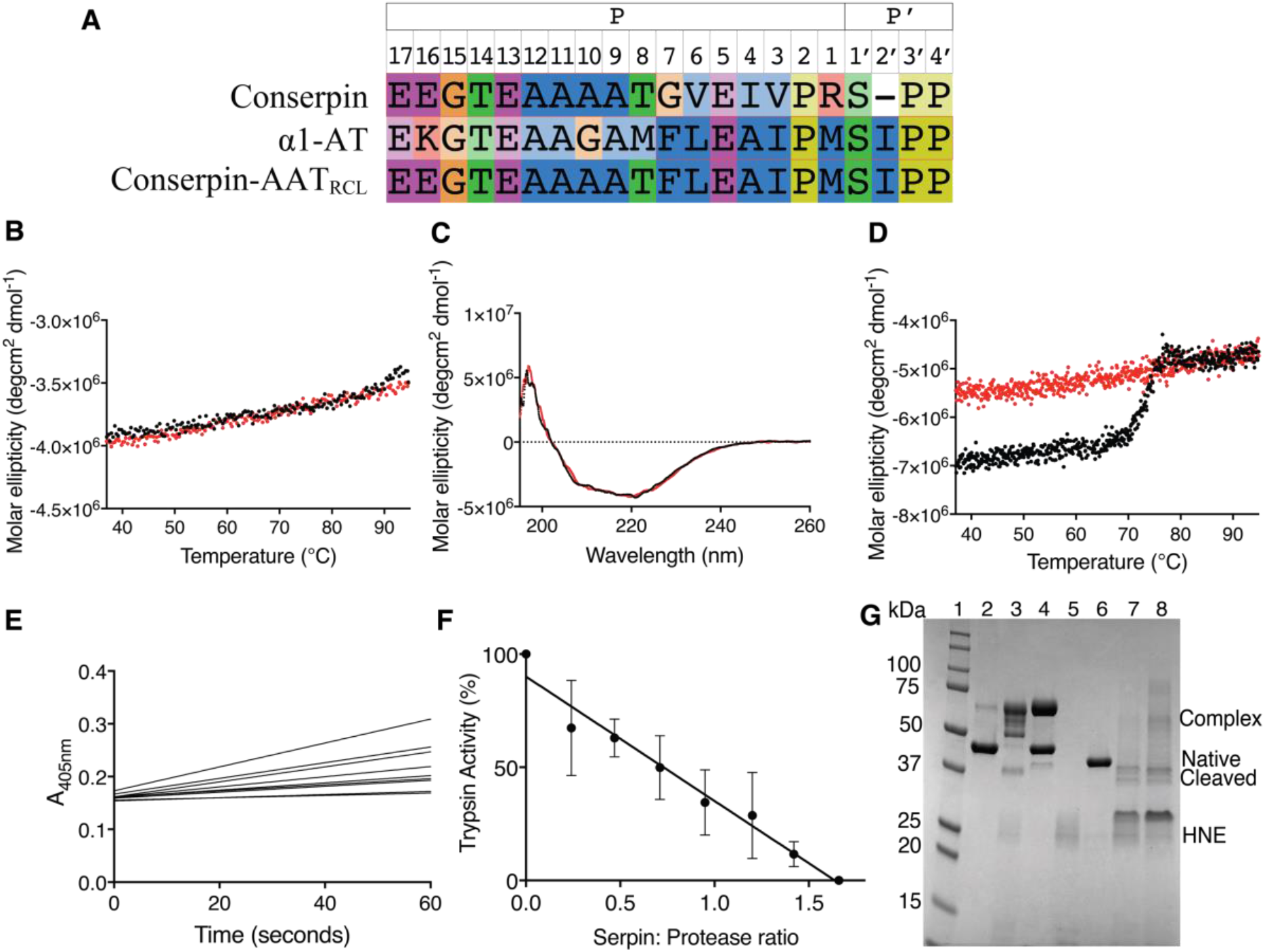
Stability and inhibitory activity of conserpin-AAT_RCL_. (A) RCL sequence alignment indicating which residues of conserpin were replaced with the corresponding residues in α1-AT; (B) Variable temperature thermal melt of conserpin-AAT_RCL_, heating to 95°C (black line) and cooling to 35°C (red line), measured by CD at 222 nm; (C) Spectral scan before (black line) and after (red line) variable temperature thermal melt; (D) Variable temperature thermal melt in the presence in 2 M GdnHCl (heating to 95°C; black line, cooling: red line); (E) Inhibitory activity assay and (F) SI against trypsin (n=3); (G) SDS-PAGE showing a serpin:protease complex formed between HNE and AAT, but less complex formed between HNE and conserpin-AAT_RCL_. From left to right: 1. Molecular weight markers (kDa); 2. α1-AT alone; 3. 1:1 ratio of α1-AT: HNE; 4. 2:1 ratio of α1-AT:HNE; 5. HNE alone; 6. conserpin-AAT_RCL_ alone; 7. 1:1 ratio of conserpin-AAT_RCL_:HNE; 8. 2:1 ratio of conserpin-AAT_RCL_:HNE.

We have previously shown conserpin to be a poor inhibitor of trypsin in comparison to α1-AT (SI = 1.8 vs 1.0 respectively)^29^. Engineering the RCL sequence of α1-AT into conserpin improves the SI against trypsin from 1.8 to 1.64 (conserpin-AAT_RCL_ SI=1.64 ± 0.2 n=3; Fig. 1E, F). Conserpin-AAT_RCL_, like conserpin, remained active after refolding upon denaturing in 6 M GuHCl (SI=2.0). Furthermore, conserpin-AAT_RCL_ does not inhibit HNE, the protease target of α1-AT. An SI could not be calculated, as there was residual HNE activity after 30-minute incubation, even with at a 2:1 serpin:protease molar ratio.

If the inhibitory pathway of serpin proceeds faster than the substrate pathway, then the SI will be close to 1. If, however, the inhibitory mechanism is too slow and the substrate pathway occurs, the SI is greater than 1^34^. SDS-PAGE using 1:1 and 2:1 serpin: protease molar ratios reveals a faint complex between conserpin-AAT_RCL_ and HNE, but also showed a large amount of cleaved species compared to the complex formation between α1-AT and HNE (Fig. 1G). Since we observe that conserpin-AAT_RCL_ is able to inhibit trypsin, and is still able to transition to the latent state upon heating, we hypothesized that the RCL mutations do not prevent its insertion into the A-sheet. We therefore sought to investigate the structure and dynamics of conserpin-AAT_RCL_ in order to identify other factors contributing to its inability to inhibit HNE.

### The role of electrostatics in the formation of a serpin:protease complex

To understand if there are any structural changes caused by modifying the RCL, we determined the X-ray crystal structure of conserpin-AAT_RCL_ in the native state (Table S1). The overall structure of conserpin-AAT_RCL_ is identical to that of conserpin—a structural alignment reveals a root mean square deviation (RMSD) of 0.2 Å across all Cα atoms. Like conserpin and indeed many other serpins, the RCL of conserpin-AAT_RCL_ is too flexible to be modelled into the electron density. Therefore, all further analyses were performed with the RCL modelled using the structure of wildtype α1-AT (PDB ID 3NE4).

Effective serpin inhibition of a protease must involve association to form an encounter complex followed by formation of a stereospecific, high-affinity complex that positions the RCL of the serpin to engage with the protease active site. Given the failure to engineer the RCL for α1-AT specificity and inhibition, we reasoned that surface electrostatics may contribute to the formation and stability of a serpin:protease complex and thus protease inhibition. The electrostatic potential surfaces of conserpin, conserpin-AAT_RCL_ and α1-AT differ in several regions. Both conserpin-AAT_RCL_ and α1-AT feature a large electropositive surface centred around the loop connecting s2B and s3B (Fig. 2B, C). In conserpin-AAT_RCL_, this patch extends to encompass the D-helix, P9–P1 of the RCL, and s2C-helix H (Fig. 2E). The corresponding region on α1-AT is much smaller, covering a region under the RCL, some residues of s1B and its connecting loop to helix G, s4B and s5B (Fig. 2F).

**Figure 2.**
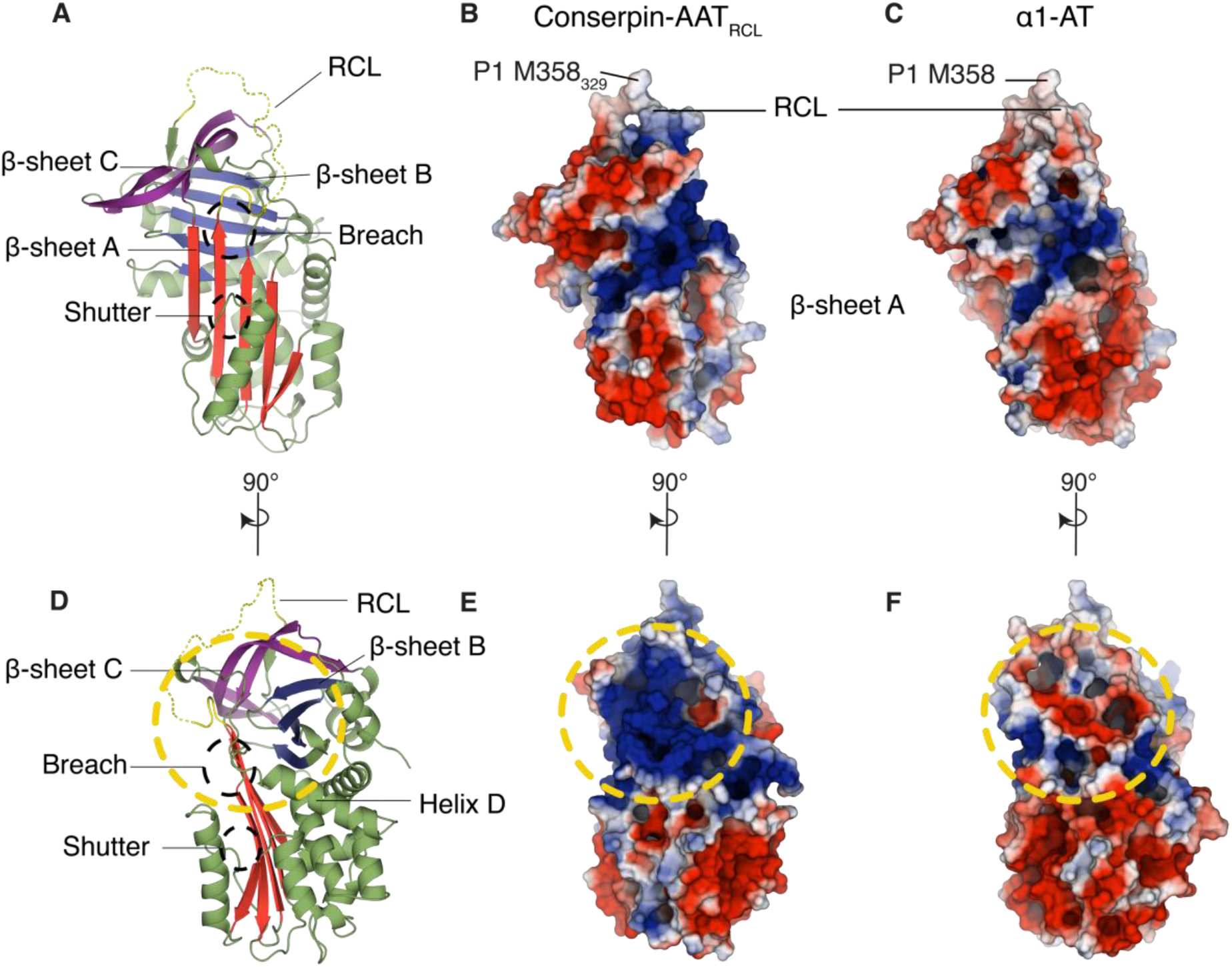
Structure and electrostatics of conserpin-AAT_RCL_. (A, D) X-ray crystal structure of native state conserpin-AAT_RCL_ represented as a cartoon. The breach and shutter regions are marked with black broken circles. (B–F) A comparison of electrostatic potential surfaces (blue=+ve, red=-ve) of (B, E) conserpin-AAT_RCL_ and (C, F) α1-AT. A large surface patch between helix D and the RCL, highlighted with yellow broken circles, has a generally positive potential in conserpin-AAT_RCL_ (E), and negative potential in α1-AT (F).

A second difference is seen on the top surface of the serpins, directly beneath the RCL. Differences between α1-AT and conserpin-AAT_RCL_—particularly in s2C, s3C, and the loop between s3A and s3C—lead to a large difference in charge on the surface beneath P9-P1 (Fig. 3A, B). In conserpin-AAT_RCL_ (and conserpin), this region has a large electropositive potential, while the corresponding region in α1-AT is more neutral in charge (Fig. 3A, B).

**Figure 3.**
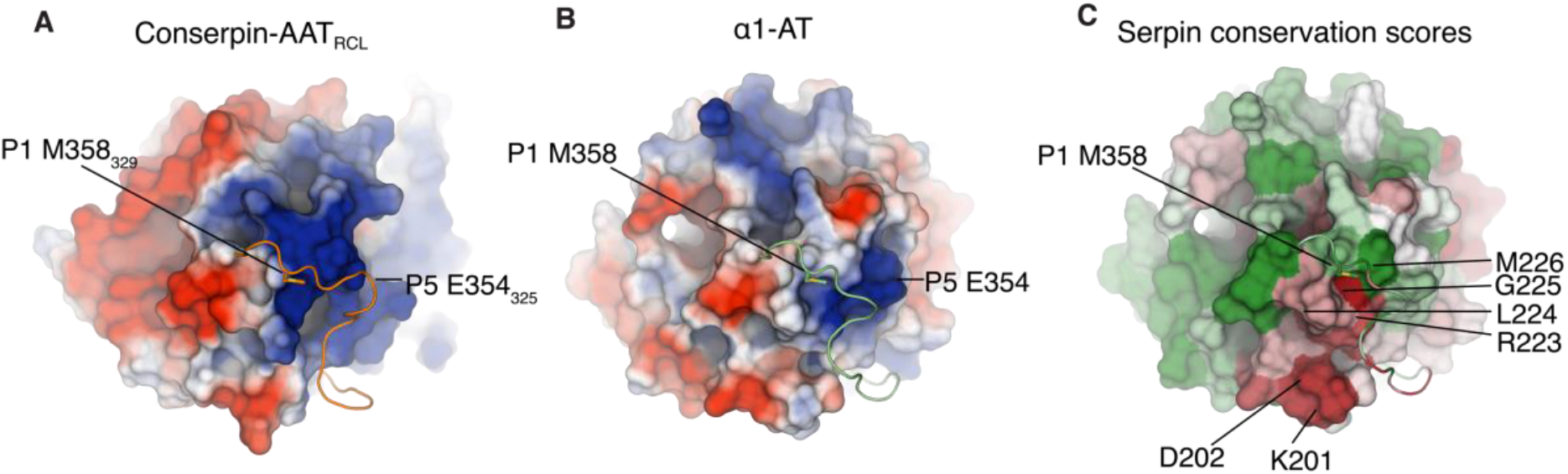
Electrostatic potential surfaces of the RCL differs between conserpin-AAT_RCL_ and α1-AT. While we have grafted the α1-AT RCL (cartoon) from P7–P2’ onto conserpin (surface), the electrostatic surface potential between conserpin-AAT_RCL_ and α1-AT differs beneath the RCL. (A) In conserpin-AAT_RCL_, the region below the RCL contains a large electropositive potential, while in α1-AT (B), the corresponding region is more neutral in charge. (C) ConSURF conservation scores for the serpin superfamily, mapped onto the surface of α1-AT as colours from forest green (highly conserved) to brick red (highly variable). This depicts poor conservation (red) of residues 201–202 and 223–225 of α1-AT, suggesting that these residues may be responsible for contributing to protease specificity within the serpin family.

Functional requirements of an inhibitory serpin’s RCL provide selective pressures on its sequence. In inhibitory serpins, the sequence of the RCL must correspond to the specificity of its target proteases^5^, maintain a linear, mobile structure in the stressed/native state, and still remain capable of insertion into highly conserved regions in β-sheet A post-cleavage (an example is the requirement of small residues in the hinge region^15,35,36^). Given these known coevolutionary pressures, it follows that there should be either highly conserved residues which are responsible for conferring this polymorphic behaviour, or a coevolutionary signal present in the sequences of functionally interacting regions within the serpin. As we were interested in the interactions between the residues of the RCL and residues beneath the RCL, we calculated conservation scores using a sequence alignment of 212 serpin sequences, and mapped them onto the structure of α1-AT (Fig. 3C). Residues facing the P1 and P1’ residues of the RCL are well conserved, compared to residues on strands s2C and s3C that face the RCL (under the residues N-terminal to P1). We were unable to identify any significant coevolutionary links between residues of the RCL and the region below it on sheet C, though this is most likely a reflection on the limited number of sequences used.

To further investigate the interactions between the RCL and the body of the serpin, we looked at the frustration networks within conserpin-AAT_RCL_ and α1-AT. In α1-AT, the RCL is minimally frustrated against the body of the serpin, with only the P12-P9 region present in a patch of high frustration. In contrast, there is a more extensive network of frustration in conserpin-AAT_RCL_, particularly between the RCL and the loop between s3A and s3C (SI Fig. 1). These distinct frustration patterns reflect the differences we observed in the electrostatics on top of the serpin body (Fig. 3), and suggest that the electrostatic compatibility between the body of the serpin and the RCL plays a key role in determining serpin functionality.

Having established clear differences in the surface electrostatics, we next investigated possible consequences for engagement with proteases trypsin and HNE. Given contrasting inhibition of these two proteases we compared their electrostatic potential surfaces. The largest difference between the two proteases is found at the active site. Whereas both proteases feature an electronegative potential in the active site cleft, in trypsin it is more extensive, encompassing S2-S4 binding pockets and the surrounding residues (Fig. 4B, 4E). In contrast, the S3-S4 binding pockets and surrounding residues of HNE contains an adjacent large electropositive patch (Fig. 4E). To observe any electrostatic potential clashes during a hypothetical serpin-protease encounter complex, we modelled a conserpin-AAT_RCL_: trypsin complex, and a conserpin-AAT_RCL_: HNE complex, each with P1 M358_329_ in the protease active site (Fig. 4A, D). The electrostatic potential for each protease and serpin were calculated separately, eliminating the influence of one electrostatic potential onto the other.

**Figure 4.**
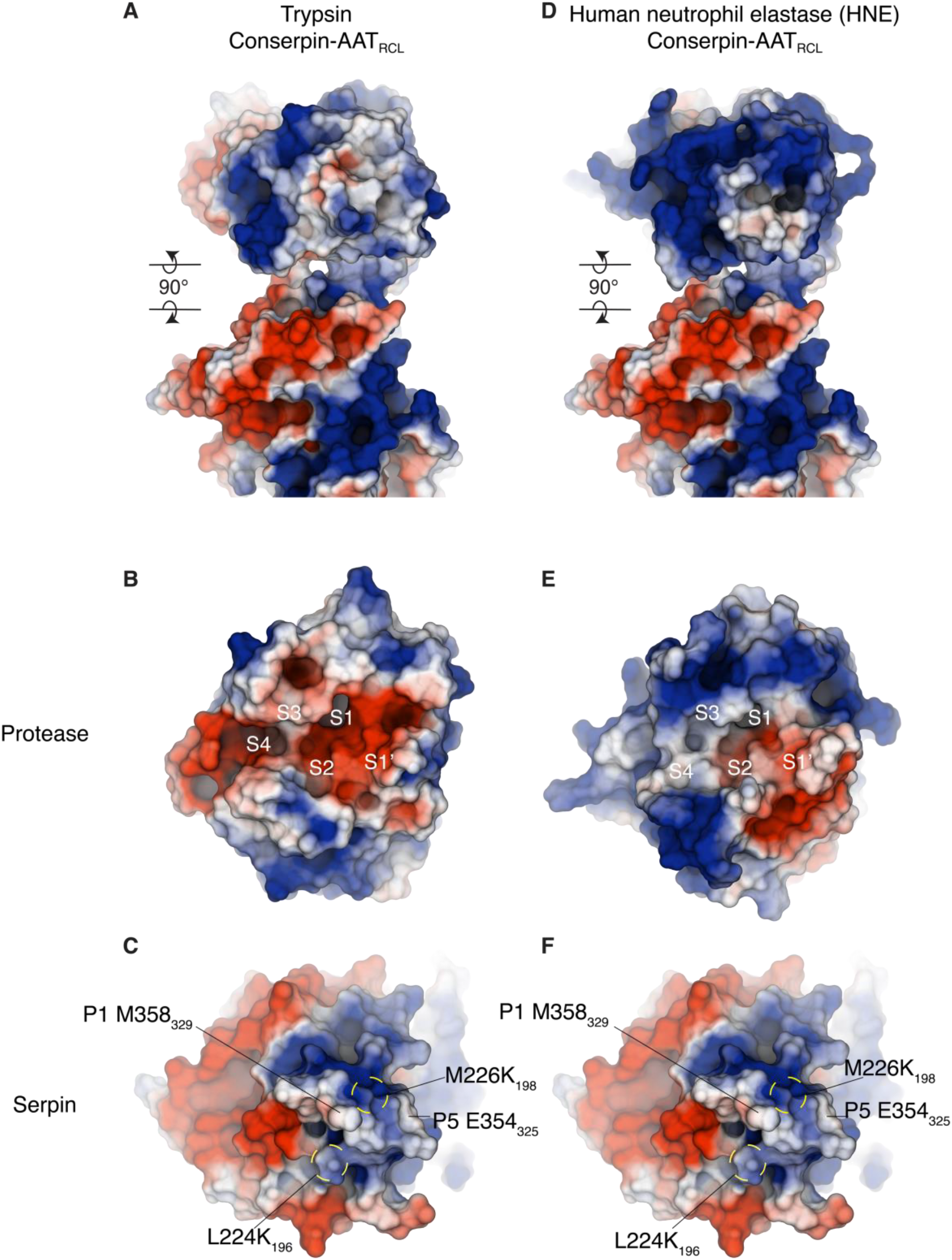
Electrostatic compatibility between serpin and protease. (A) Electrostatic surfaces of a modeled complex between trypsin and conserpin-AAT_RCL_, and (D) between HNE and conserpin-AAT_RCL_. Associated complexes are separated into individual proteins by rotating each molecule by 90° around the horizontal axis in the plane of the paper (clockwise for the top molecules, anti-clockwise for the bottom molecules). (B) Electrostatic surface for the active site of trypsin and (E) HNE shows that trypsin has a more electronegative binding cleft than HNE. Comparing this to the electrostatic surface of (C, F idem.) conserpin-AAT_RCL_ suggests a greater electrostatic compatibility between trypsin, particularly the electropositive surface below the RCL. However, the electropositive surface of S3-S4 binding pocket in HNE suggests there may be a charge repulsion with the electropositive surface potential of conserpin-AAT_RCL_ at P6–P3.

The calculated electrostatic potential suggests trypsin has greater electrostatic compatibility with conserpin-AAT_RCL_ than HNE. This compatibility can be attributed to the large electropositive surface of the RCL and the body below the RCL. Compatibility will be essential for the formation and stability of a Michaelis serpin-protease complex, where there is contact between the binding pockets (S4-S1’) of the protease and P6-P1’ residues of the RCL. The formation and stability of a serpin-protease complex between conserpin-AAT_RCL_ and trypsin can occur with favourable interaction between trypsin’s electronegative S3-S4 pockets (Fig. 4B) and the electropositive potential of conserpin-AAT_RCL_ P6-P3 residues (Fig. 4C). Therefore, conserpin-AAT_RCL_ can inhibit trypsin. In comparison, the formation and stability of a complex may be hindered by the charge-charge repulsion between the electropositive S3-S4 binding pockets of HNE (Fig. 4D) and the electropositive surface of P6-P3 of the RCL (Fig. 4E). As a result, conserpin-AAT_RCL_ behaves as a substrate to HNE rather than as an inhibitor.

### RCL dynamics are important for protease inhibition

Given the large conformational changes involved in serpin function, and specifically the central role played by the RCL in protease engagement and subsequent insertion into the A-sheet, an investigation of the dynamics of the RCL of conserpin-AAT_RCL_ may provide some insight into its inhibitory properties. We therefore performed molecular dynamics (MD) simulations of conserpin-AAT_RCL_ and compared the results to those of α1-AT and conserpin simulations we performed previously^29^. Although we are unable to perform simulations for long enough to observe the RCL insertion into the A-sheet, MD is able to reveal the intrinsic dynamics of the RCL and specifically the lifetime of its interactions with the body of the serpin. After reaching equilibrium at around 150 ns, the root mean square deviation (RMSD) indicated that the simulations remained stable with no large conformational changes observed. Given the importance of RCL conformation in facilitating the S→R transition following protease engagement, we analyzed the dynamics of the A-sheet and the RCL, and also the interactions between the RCL and the body of the serpin during the time course of the MD simulations. The central A-sheet contains two conserved regions, the shutter and breach, which are critical for the insertion of the RCL; mutations in these regions often render the serpin susceptible to misfolding and aggregation^37^.

Substitution of part of the conserpin RCL for the corresponding region of α1-AT did *not* serve to reduce the flexibility of the RCL region to the lower level observed in α1-AT. Instead, while the region of conserpin-AAT_RCL_ around residues 353_324_-362_333_ showed a reduced root mean square fluctuation (RMSF) (corresponding to less conformational variability), the region around residues 342_314_-352_323_ showed an increased RMSF (Fig. 5D). This increase in flexibility is evident in comparing MD snapshots of the three systems, where the substantially increased flexibility of the lower RCL region of conserpin-AAT_RCL_ is clearly evident (Fig. 5A-C). This highly dynamic region encompasses the hinge region of the RCL, the first residues that insert into the A-sheet.

As the fate of the serpin as either a substrate or inhibitor is determined by the competition between rates of RCL insertion and the de-acylation of the protease^14^, it is likely that an increase in RCL loop dynamics would slow the rate of insertion. This would allow a protease with a fast catalytic mechanism (such as HNE), to escape inhibition, thereby pushing the serpin down the substrate pathway.

**Figure 5.**
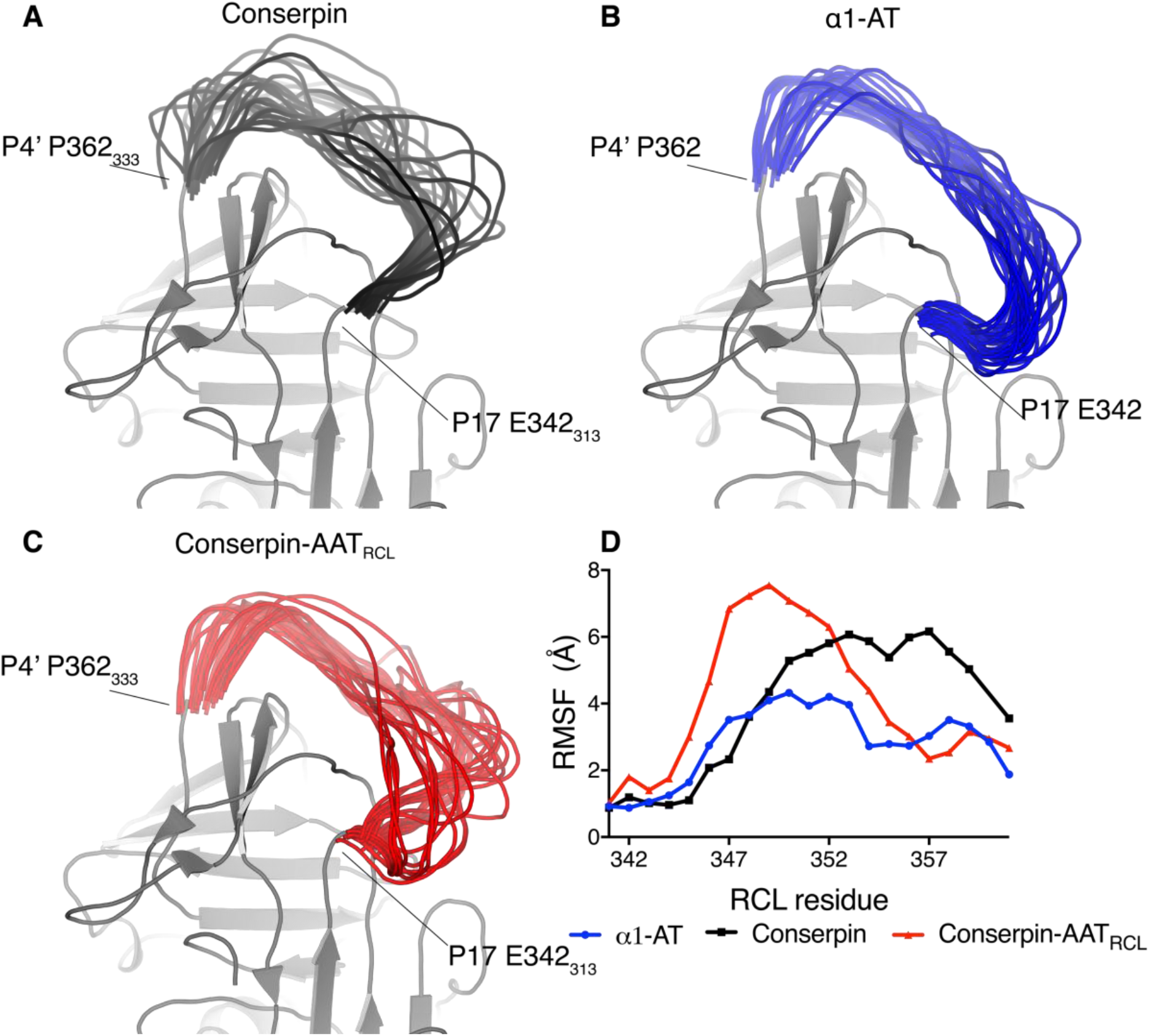
The dynamics of the RCL is important for inhibition. Snapshots of conformations of the RCL from the MD runs at 50 ns intervals overlaid on static structure for the rest of the molecule, showing that (A) conserpin prefers an extended-hinge RCL conformation, (B) α1-AT prefers a bent-hinge RCL conformation, and (C) conserpin-AAT_RCL_ occupies both of these conformations. the increased flexibility of the lower RCL region (residues 342_314_-352_323_) relative to both conserpin and α1-AT. (D) Root mean square fluctuation (RMSF) calculated for the RCL region from the molecular dynamics simulations shows that the conserpin-AAT_RCL_ (red) has lower flexibility than conserpin (black) in the 3 5 3 324-3 6 2333 region but a higher flexibility in the 342314-352323 region than conserpin and α1-AT (blue) (α1-AT numbering), reflecting the structural differences between the two conformational clusters occupied by the RCL.

To understand the RCL conformations adopted by conserpin-AAT_RCL_ throughout the simulations, we performed principal component analysis on the conformations of the RCL backbone (between P17-P1’) over all simulations (α1-AT, conserpin and conserpin-AAT_RCL_), followed by a clustering. This produced a total of 9 clusters, with RCL conformations within each cluster being structurally close but clearly distinguishable from others (SI Fig. 2). These analyses show that α1-AT’s RCL maintains a reasonably close set of conformations throughout the three independent simulations, while conserpin’s RCL explores a broad variety of conformations that are exclusive of those explored by α1-AT’s RCL (Fig. 6A). Conserpin-AAT_RCL_’s RCL not only adopts conformations that overlap with those of the other serpins, but also explores conformations that were not seen in α1-AT and conserpin simulations.

**Figure 6:**
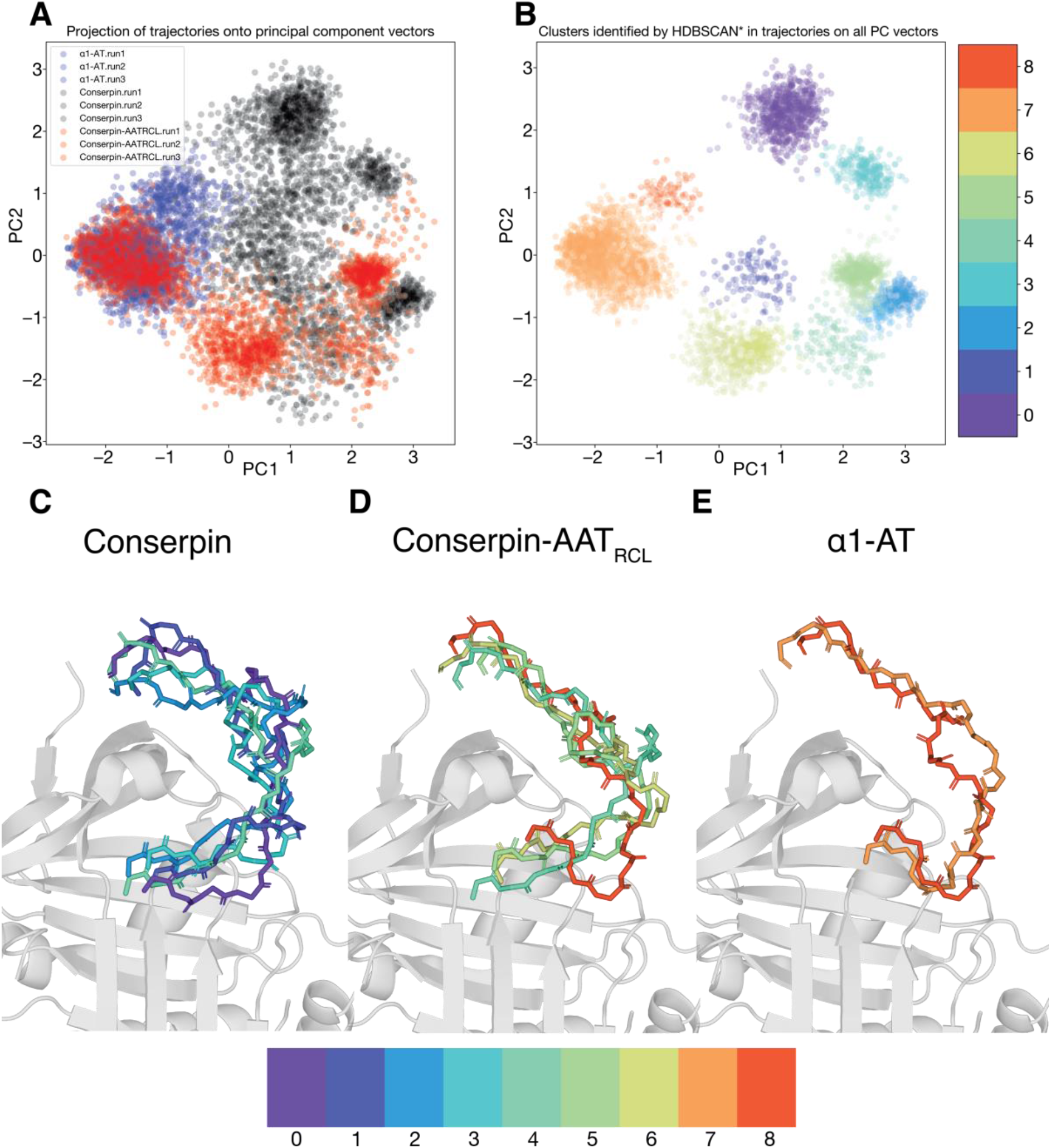
RCL conformational cluster determination by principal component analysis. To describe the motion of the RCL across all simulations, principal component vectors were determined for all RCL backbone conformations. (A) The trajectories of each RCL (α1-AT: blue, conserpin: black, conserpin-AAT_RCL_: red) are projected on the first 2 PC axes, and (B) these conformations were grouped into 9 clusters. For (C) conserpin, (D) conserpin-AAT_RCL_ and (E) α1-AT, representative RCL backbone conformations for the clusters explored by each serpin over the course of the molecular dynamics simulations, are shown atop a serpin body (grey cartoon α1-AT (PDB: 3NE4)).

Conserpin’s RCL explored 5 different conformations (5 clusters), most of which have an extended conformation in which the hinge region of the RCL is moved away from the breach region of β-sheet A (Fig. 6B). This preference for an extended RCL hinge in conserpin is surprising, as conserpin has an extended salt bridge network in the breach region in comparison to α1-AT^29^, which was hypothesised to stabilise conserpin’s native state.

For α1-AT, the RCL explores 2 similar conformations, with both conformations containing the hinge region primed for insertion between s3A and s5A. This expands on the RMSF analysis (Fig. 5D), where α1-AT’s RCL was seen to be relatively rigid over the course of the simulations (in comparison to conserpin and conserpin-AAT_RCL_). The rigidity of α1-AT’s RCL suggests that there are interactions between the body and the RCL that reduce the dynamics of the loop, and possibly prime the hinge region between s3A and s5A strands.

The RCL of conserpin-AAT_RCL_ explores 4 conformations: one overlapping with a cluster seen in conserpin, one overlapping with a cluster seen in α1-AT, and 2 conformations unique to conserpin-AAT_RCL_. All of these conformations (except the α1-AT-like one) include an extended hinge region away from β-sheet A. This could possibly be a consequence of the interactions between the residues on β-sheet C and the RCL, as stated previously^38^. Interestingly, one of the conformations include a slight helical turn from P10-P7 (similarly to an α1-AT / α1-antichymotrypsin chimera^39^), possibly responsible for pulling the hinge region away from β-sheet A. Importantly, one of conserpin-AAT_RCL_’s RCL conformation is similar to α1-AT’s, where the hinge region is primed to insert into β-sheet A. The ability of conserpin-AAT_RCL_ to access this conformation may explain the increase in inhibitory activity against trypsin over conserpin, as the conserpin-AAT_RCL_ RCL could insert faster than conserpin from this pose. However, despite this primed hinge region conformation, conserpin-AAT_RCL_ primarily remains a substrate against HNE. It is possible that HNE could negatively impact on the conformation of the RCL upon encounter, or even prevent formation of a stable serpin:protease complex, a scenario in which HNE’s catalytic mechanism occurs more rapidly than trypsin, allowing for rapid cleavage of the RCL followed by substrate rather than inhibitor behaviour. A structural difference was also observed by calculating the phi-psi angles of the RCL for each serpin over the course of the simulations. Replacement of P7-P2’ of α1-AT onto conserpin has failed to reproduce the conformational pattern seen in α1-AT. Specifically, while the conformations adopted by conserpin-AAT_RCL_ in the region around residues 353324-362333 (SI Fig. 3, red) are more similar to those of α1-AT than conserpin (SI Fig. 3, blue and black, respectively), those in the region around residues 342_314_-352_323_ (SI Fig. 3) show a distinctly different set of conformations. Together these observations indicate that the conformational landscape sampled by the structure in and around the RCL region, including areas near the breach region, is important in the process of RCL insertion, and thus ultimately serpin inhibitory specificity.

## Discussion

Conserpin shares high sequence identity to α1-AT (59%), is extremely stable, polymerisation-resistant and yields large quantities when expressed through recombinant techniques. It is therefore an attractive model system for investigating the folding, stability and function of serpins. In this study, to investigate the determinants of serpin specificity, we used a conserpin/α1-AT chimera, which we call conserpin-AAT_RCL_, in which the residues in the RCL are replaced with those of α1-AT. The resulting hybrid retained the thermostability and polymerisation resistance of conserpin. However, despite containing the RCL sequence of α1-AT, which is thought to be a key determinant of inhibitory specificity, conserpin-AAT_RCL_ showed only minor improvement as an inhibitor of trypsin, in comparison to conserpin, and like conserpin behaved mostly as a substrate against HNE.

We attempted to rationalise the substrate behaviour of conserpin-AAT_RCL_ using a structural and molecular modelling/simulation approach. Although the X-ray crystal structure of conserpin-AAT_RCL_ revealed no significant differences with the parent molecule, we were able to provide insights into the failure to transfer specificity by analysing electrostatic differences and changes in the flexibility of RCL, hinge, breach and shutter regions with molecular dynamics simulations.

For a serpin to carry out its inhibitory function, the serpin and protease must come into contact with each other. Reasoning that, like other protein-protein complexes^40,41^, cognate serpins and proteases must exhibit complementary electrostatic surfaces to ensure rapid, and high affinity association, we identified several differences between the electrostatic surface characteristics of α1-AT and conserpin-AAT_RCL_ that may contribute to their contrasting inhibitory properties. HNE contains a shallow active site that interacts with P6-P3’ residues of the RCL, therefore the electrostatic surface of this region must be complementary to ensure efficient binding to the P1 methionine residue^25,42^. In comparison to α1-AT, conserpin-AAT_RCL_ harbours several regions where poor charge complementarity may explain the diminished capacity to form a complex with HNE, and subsequently why it acts as a substrate rather than an inhibitor. One of these regions includes the electrostatic potential beneath the RCL. The role of electrostatics has been investigated for several serpins. For example, single-pair Förster resonance energy transfer (spFRET) studies of the inhibition of anionic rat and cationic bovine trypsin by α1-AT showed only partial translocation of anionic rat trypsin compared to full translocation of cationic bovine trypsin^43,44^. This indicates that the electrostatic potential between the protease and serpin are important for formation of a serpin:protease complex and protease inhibition. Similarly, for the serpin PAI-1, the Michaelis complex between tissue-type plasminogen activator (tPA) and PAI-1 was observed to have more complementary electrostatic interactions than the complex between urokinase-type plasminogen activator (uPA) and PAI-1. This was used to explain the difference in second-order inhibitory rate constants between the two proteases: tPA is inhibited at a faster rate (2.6 × 10^7^ M^-1^ s^-1^) compared to uPA (4.8 × 10^6^ M^-1^ s^-1^) ^45,46^. Furthermore, an arginine to glutamic acid substitution produced a ‘ serpin-resistant’ tPA variant, where the glutamic acid produced a repulsion to PAI-1, leading to a failure to inhibit this variant. This tPA variant was only inhibited through creating a complementary PAI-1 with the opposite glutamic acid to arginine mutation^47,48^, further emphasising the importance of surface potential in the formation of a stable serpin:protease complex. Any possible repulsive interactions may destabilize a serpin:protease complex and therefore prevent inhibition.

Dynamics in the RCL is important for its insertion into β-sheet A during protease inhibition. We therefore investigated the difference in RCL dynamics between conserpin-AAT_RCL_ and α1 - AT using molecular dynamics simulations. Despite P7-P2’ of conserpin-AAT_RCL_ being identical to the corresponding region in α1-AT, the overall flexibility of the RCL region as a whole was not reduced to the level of α1-AT, while the hinge region of the RCL, which inserts into β-sheet A first during insertion, exhibited higher flexibility in conserpin-AAT_RCL_ compared to α1-AT. A plausible explanation for this is the additional residue at P2’. Conserpin was designed without an isoleucine at P2’, producing an RCL length that fits onto the serpin body. The addition of P2’ isoleucine in conserpin-AAT_RCL_ may force the RCL to adopt a non-ideal conformation, possibly increasing the dynamics of the hinge region.

The conformation of the RCL is likely highly tailored to the particular inhibitory specificity of each serpin. α1 -AT, a potent inhibitor of HNE, has an RCL that is in a primed position for insertion into β-sheet A. That is, the hinge region is poised between strands 3A and 5A, allowing for rapid insertion during HNE inhibition. Conserpin and conserpin-AAT_RCL_ contain RCL conformations that are extended, with the hinge region away from β-sheet A, likely reducing the rate at which the RCL can insert. One conformation that conserpin-AAT_RCL_ explores contains a primed hinge region, which possibly explains the increase in its inhibitory activity against trypsin (compared to conserpin), but is not enough to produce inhibition against HNE. Furthermore, the breach region of α1-AT ‘loosens’ and opens over the course of the simulations^29^, while the breach region in conserpin and conserpin-AAT_RCL_ remains rigid due to the extended salt bridge network. Therefore, it is possible that α1-AT inhibits HNE at a rapid rate due to the primed position of the hinge region and opening of the breach region, allowing inhibition before HNE’s de-acylation step of cleavage. Our observation that the RCL of conserpin-AAT_RCL_ sampled this primed hinge conformation out of 4 possible conformations, and the relatively rigid nature of its breach region, suggests that the initial steps of RCL insertion into A-sheet are slower than the de-acylation step of HNE’s cleavage.

Previous studies that have attempted to convert the specificity of ACT to that of α1-AT by swapping RCL residues have been generally unsuccessful, as the chimeras had a greater SI and slower inhibitory rate compared to α1-AT. This suggests that other factors may play important roles, including interactions between the RCL and the body of the serpin, and the structure of the chimeric RCL^13,24,25,39^. Engineering of ACT/α1-AT chimeras show that HNE’s proteolytic mechanism occurs on a shorter timescale in comparison to ACT’s catalytic mechanism^13^. It is also known that an increase in the dynamics of the RCL can affect the serpin’s ability to inhibit a protease. Notably, loss of a salt bridge in the breach region in α1-AT Z variant increases RCL dynamics and subsequently leads to an SI increase (from 1.0 to 1.8) and decrease in rate of inhibition (from 6.9 to 2.3 × 10^6^ M^-1^ s^-1^)^49–52^. This implies that the rate of RCL insertion occurs slower than the de-acylation step of HNE’s catalytic mechanism, producing a substrate rather than an inhibitor of HNE. It is likely that RCL-protease interactions will vary for each protease, influencing the dynamics of the RCL^27^ and the conformational change needed for RCL insertion^53^. With the use of fluorescent labels, it was observed that the two protease targets of plasminogen activator inhibitor-1 (PAI-1), tissue-type plasminogen activator (tPA) and urokinase-type plasminogen activator (uPA), rests differently on the P1-P1’ bond and change the dynamics of the RCL when bound. tPA affects the C-terminus of the RCL through exosite interactions, while retaining dynamics observed with free PAI-1. In contrast, uPA affects the N-terminus with different exosite interactions, restricting the dynamics and immobilising the RCL. This difference in RCL dynamics also contributes to the difference in the rates the proteases are inhibited by PAI-1^27^. Taken together, the dynamics of the RCL is critical for the rate of insertion during protease inhibition. Fast insertion favours protease inhibition while slow insertion forces the serpin to undergo the substrate pathway.

Along with the possibility of electrostatic repulsion and increased RCL dynamics, the failure to transfer specificity onto conserpin-AAT_RCL_ could be a consequence of the delicate balance between stability and function. Serpins use the metastable conformation to undergo the large conformational change necessary for its inhibitory function. Increasing the stability of this metastable state may decrease the dynamics and plasticity required to undergo the S→R transition during inhibition of a target protease. For example, increasing the stability of α1-AT more than 13 kcal mol^-1^ than the wild type α1-AT compromises its inhibitory activity^54^. For conserpin, the very high stability, although still functional, can be attributed to certain key regions important for the serpin’s inhibitory mechanism and S→R transition^29^. Structural plasticity is required in the breach region, as this region is important in controlling the insertion of the RCL and conformational change to allow for protease inhibition. The extensive salt-bridge network in the breach region in conserpin and conserpin-AAT_RCL_ increases rigidity and slows the opening of β-sheet A between strands s3A and s5A. The rigidity of the breach, along with displacement of conserpin and conserpin-AAT_RCL_ hinge region away from β-sheet A, may explain the reduced inhibitory activity of conserpin towards trypsin, and the failure of conserpin-AAT_RCL_ to inhibit HNE. Furthermore, helix F, which packs tightly against the A-sheet, may act as a barrier to RCL insertion via A-sheet opening, and must partially unfold to allow rapid RCL insertion^55–57^. Mutations on the helix F/A-sheet interface of α1-AT can relieve this tight packing, increasing the stability but also decreasing activity. Therefore, the tight packing between helix F and A-sheet contributes to the metastability and that is relieved in the S→R transition. In conserpin, this interface is tightly packed, but not to the extent of α1-AT. As a result, the tight packing at this interface may not have the strain observed in α1-AT, slowing the partial unfolding of helix-F to allow for rapid RCL insertion.

## Conclusions

In this work, we utilized a serpin chimera to investigate the rules that govern serpin specificity, by studying the effect of replacing the RCL of conserpin, a model synthetic serpin, with the corresponding sequence from α1-AT. Despite possessing the RCL sequence of α1-AT, specificity against trypsin or HNE was not restored to that of α1 - AT. Crystal structural analysis and molecular dynamics simulations indicate that, although the RCL sequence may partially dictate specificity, electrostatic surface potential coupled with dynamics in and around the RCL likely play an important role. The dynamics of the RCL appears to govern the rate of insertion during protease inhibition, dictating whether it behaves as an inhibitor or a substrate. The unusual mechanism of serpin action also requires a delicate balance between stability, dynamics and function. Engineering serpin specificity is therefore substantially more complex than solely manipulating the RCL sequence, and although may be guided by the general principles discussed in this work, each serpin will most likely present unique challenges. Notwithstanding this, further characterisation of the role of dynamics will be required to advance our understanding of how serpins perform their exquisite inhibitory functions.

## Methods

### Design of conserpin-AAT_RCL_

The design of conserpin-AAT_RCL_ was based of the RCL sequence of α1-AT. Residues P7-P2’ of the α1-AT RCL were mutated onto the original conserpin molecule to provide specificity against trypsin and neutrophil elastase. The residue numbering adheres to that adopted as previous^29^: Q105α1-AT and corresponding conserpin-AAT_RCL_ residue R79conserpin-AAT_RCL_ is written as Q105R79.

### Expression constructs

The plasmid encoding conserpin-AAT_RCL_ was generated using ligation-independent cloning with the pLIC-HIS vector^58^ using standard protocols; this construct was transformed into BL21(DE3) pLysS E. coli.

### Protein expression and purification

Protein was expressed 2xYT media and induced with isopropyl ß-D-1-thiogalactopyranoside (IPTG) at an OD600 of 1. Expression was continued for 3 hours before cells were harvested and lysed in 10 mM imidazole, 50 mM NaH2PO4, 300 mM NaCl, pH 8.0. Following centrifugation, the soluble fraction was applied to nickel-NTA loose resin (Qiagen) and washed with 50 mL of 20 mM imidazole, 50 mM NaH2PO4, 300 mM NaCl, pH 8.0. Bound protein was eluted with 250 mM imidazole, 50 mM NaH2PO4, 300 mM NaCl, pH 8.0. Fractions containing protein were loaded into a Superdex 200 16/60 column for further purification and eluted with 50 mM tris-HCl, 150 mM NaCl pH 8.0.

### Characterisation of inhibitory properties

The stoichiometry of inhibition against bovine trypsin (Sigma-Aldrich) was performed similarly as described^29,34^. Briefly, various concentrations of conserpin-AAT_RCL_ was incubated with a constant trypsin concentration at 37°C for 30 min in 50 mM tris-HCl, 150 mM NaCl, 0.2% v/v PEG 8000 pH 8.0. The residual trypsin activity was measured at 405 nm using the substrate Na-benzoyl-L-arginine 4-nitroanilide hydrochloride (Sigma-Aldrich).

To test for activity after refolding, conserpin-AAT_RCL_ was unfolded in 6 M guanidine hydrochloride (GndHCl) 50 mM tris-HCl, 150 mM NaCl pH 8.0 for 2 hours before refolding via dilution for another 2 hours. Any aggregate was pelleted by centrifugation and the sample dialysed against the same buffer to remove any remaining GndHCl. The SI assay against trypsin was performed as stated.

To observe an SDS-stable serpin: protease complex, different ratios of serpin were incubated with protease for 30 minutes at 37°C. SDS reducing dye was added to each sample and quenched on ice to stop any further reaction. Samples were loaded onto a 10% SDS-PAGE.

### Circular dichroism scans and thermal denaturation

Circular dichroism (CD) measurements were performed on a Jasco J-815 CD spectrometer at a protein concentration of 0.2 mg/mL with PBS using a quartz cell with a path-length of 0.1 cm. Far-UV scans were performed at 190–250 nm. For thermal denaturation, a heating rate of 1°C/min from 35°C to 95°C was used, with the change in signal measured at 222 nm. For samples containing 2 M GdnHCl, refolding was measured directly after the thermal melt by holding the temperature at 95°C for 1 min before the temperature was decreased to 35°C at the same rate. The midpoint of transition (Tm) was obtained by fitting the data with a Boltzmann sigmoidal curve in accordance with the method described^29^ for both forward and reverse thermal denaturation experiments.

### Crystallization, X-ray data collection, structure determination and refinement

Crystals of conserpin-AAT_RCL_ were obtained using hanging drop vapour diffusion, with 1: 1 (v/v) ratio of protein to mother liquor (total well volume of 500 μL). The protein was concentrated to 10 mg/mL and crystals appeared in 0.2 M magnesium chloride, 0.1 mM Bis-Tris and 20% PEG 3350, pH 6.5 after 5 days.

Diffraction data was collected on the MX2 beamline at the Australian Synchrotron. The diffraction data was processed with iMOSFLM^59^ to 2.48 Å, followed by scaling with SCALA^60^ in the CCP4 suite^61^. The structure was determined by molecular replacement (MR) with Phaser^62^, using conserpin (native state) structure as a search probe (PDB 5CDX)^29^. The model was built and refined using PHENIX^63^ and Coot^64^.

### Computational resources

Atomistic MD simulations were performed on Multi-modal Australian ScienceS Imaging and Visualisation Environment (MASSIVE), and in-house hardware (NVIDIA TITAN X Pascal GPU).

### Atomic coordinates, modelling and graphics

The RCL was modelled onto the x-ray crystal structure using MODELLER^65^. In MD simulations, atomic coordinates were obtained from the following PDB entry: 3NE4^66^. α1-AT and conserpin molecular dynamics simulations used for the analysis were run previously in the original Conserpin paper^29^. The residue numbering remained as determined by crystal structure, that is, the glutamine from the TEV cleavage tag remained as residue-1. Structural representations were produced using PyMOL version 2.0.4^67^ and VMD 1.9.4^68^. Trajectory manipulation and analysis was performed using MDTraj^69^ and VMD 1.9.4^68^. Electrostatic calculations were performed with the APBS plugin^70,71^ on PyMOL. Serpin:protease complexes were modelled based on the X-ray crystal structure of a serpin:protease Michaelis complex (*Manduca sexta* serpin 1B with rat trypsin (S195A), PDB: 1K9O)^72^.

### Molecular dynamics (MD) systems setup and simulation

Each protein, with protonation states appropriate for pH 7.0 as determined by PROPKA^73,74^, was placed in a rectangular box with a border of at least 10 Å, explicitly solvated with TIP3P water^75^, counter-ions added, and parameterized using the AMBER ff99SB all-atom force field^76–78^. After an energy minimization stage and an equilibration stage, production simulations were performed in the NPT ensemble using periodic boundary conditions and a time step of 2 fs. Temperature was maintained at 300 K using the Langevin thermostat. Three independent replicates of each system were simulated for 500 ns each using Amber 14^79^. The three independent replicates for each system were concatenated, and RMSD, RMSF, and phi and psi angles computed over 500 ps timesteps using VMD 1.9.4^68^.

### Sequence methods

In calculating construct sequence identities, construct sequences were aligned using MUSCLE^80,81^ v3.8.1551. The 6xHIS-TEV-SacII N-terminal peptide was removed from the alignment so as not to inflate alignment statistics. Percentage identities were calculated as %id = 100% × number of identity columns / length of aligned region (including gaps).

Mapping of sequence conservation on structure α1-AT^82^ was performed using the Consurf 2016 server^83^ using the previously designed alignment of 212 serpins^36^. Sequence coevolution analysis was performed using the OMES χ^2^ residue independence test^84^, as well as the SCA^85^ and ELSC^86^ perturbation-based residue covariance methods.

### RCL principal component analysis & clustering

From the nine trajectories described above, trajectories of the 72 atoms describing the backbone (N, CA, C, O) from P17 (E342)–P1’ (S359) were extracted using MDTraj^87^. These trajectories were concatenated together into a 8993-frame trajectory, and Scikit-learn^87^ was used to calculate eigenvectors describing 216 principal component vectors. The top three PCA vectors describe 35.64%, 16.67%, and 11.72% respectively of the variance across all conformations in the concatenated trajectory. The nine trajectories were then transformed into this PCA space, and plotted using matplotlib^88^.

The concatenated trajectory, as expressed in PCA coordinates, was clustered using the HDBSCAN algorithm^89,90^, using default parameters, except a minimum cluster size of 1% of the total trajectory (90 frames).

### Frustration calculation

Local frustration analysis of the modelled serpin: protease complexes was conducted with the Frustratometer2 web server^91^. Essentially, the energetic frustration is obtained by the comparison of the native state interactions to a set of generated “decoy” states where the identities of each residue are mutated. The constant *k* used to model the electrostatic strength of the system was set to its default value (4.l5). A contact is defined as “minimally frustrated” or “highly frustrated” upon comparison of its frustration energy with values obtained from the decoy states.

## Author contributions

EMM and AMB designed the study. EMM performed the protein expression, purification, CD thermal melt and assay experiments with assistance from BTP and DEH. EMM, SM and AMB performed the crystallography, with assistance from BTP. JF performed molecular dynamics simulations. JF and BTR analyzed simulations. MGSC performed frustration analysis. EMM and BTR generated figures. EMM, BTR, SM and AMB wrote the manuscript.

## Acknowledgements

BTP is a Medical Research Council Career Development Fellow. SM acknowledges fellowship support from the Australian Research Council (FT100100960). We thank the Australian Synchrotron for beam-time and technical assistance. This work was supported by the Multimodal Australian ScienceS Imaging and Visualisation Environment (MASSIVE) (www.massive.org.au). We acknowledge the Monash Protein Production Unit and Monash Macromolecular Crystallization Facility.

## Accession Numbers

The coordinates and structure factors have been deposited in the Protein Data Bank under accession code 6EE5.

## References

(1) Huntington, J. A., Read, R. J., and Carrell, R. W. (2000) Structure of a serpin-protease complex shows inhibition by deformation. Nature 407, 923–926.

(2) Elliott, P. R., Lomas, D. A., Carrell, R. W., and Abrahams, J. P. (1996) Inhibitory conformation of the reactive loop of alpha 1-antitrypsin. Nat. Struct. Biol. 3, 676–681.

(3) Stratikos, E., and Gettins, P. G. W. (1999) Formation of the covalent serpin-proteinase complex involves translocation of the proteinase by more than 70 A and full insertion of the reactive center loop into-sheet A. Proc. Natl. Acad. Sci. 96, 4808–4813.

(4) Tew, D. J., and Bottomley, S. P. (2001) Probing the equilibrium denaturation of the serpin alpha(1)-antitrypsin with single tryptophan mutants; evidence for structure in the urea unfolded state. J. Mol. Biol. 313, 1161–1169.

(5) Gettins, P. G. W. (2002) Serpin structure, mechanism, and function. Chem. Rev. 102, 4751–804.

(6) Krishnan, B., and Gierasch, L. M. (2011) Dynamic local unfolding in the serpin α-1 antitrypsin provides a mechanism for loop insertion and polymerization. Nat. Struct. Mol. Biol. 18, 222–6.

(7) Tsutsui, Y., Dela Cruz, R., and Wintrode, P. L. (2012) Folding mechanism of the metastable serpin α1-antitrypsin. Proc. Natl. Acad. Sci. U. S. A. 109, 4467–72.

(8) Cabrita, L. D., and Bottomley, S. P. (2004) How do proteins avoid becoming too stable? Biophysical studies into metastable proteins. Eur. Biophys. J. 33, 83–88.

(9) James, E. L., and Bottomley, S. P. (1998) The mechanism of alpha 1-antitrypsin polymerization probed by fluorescence spectroscopy. Arch. Biochem. Biophys. 356, 296–300.

(10) Dupont, D. M., Madsen, J. B., Kristensen, T., Bodker, J. S., Blouse, G. E., Wind, T., and Andreasen, P. A. (2009) Biochemical properties of plasminogen activator inhibitor-1. Front. Biosci. (Landmark Ed. 14, 1337–61.

(11) Mushunje, A., Evans, G., Brennan, S. O., Carrell, R. W., and Zhou, A. (2004) Latent antithrombin and its detection, formation and turnover in the circulation. J. Thromb. Haemost. 2, 2170–2177.

(12) Padrines, M., Schneider-Pozzer, M., and Bieth, J. G. (1989) Inhibition of neutrophil elastase by alpha-1-proteinase inhibitor oxidized by activated neutrophils. Am. Rev. Respir. Dis. 139, 783–90.

(13) Rubin, H., Plotnick, M., Wang, Z. M., Liu, X., Zhong, Q., Schechter, N. M., and Cooperman, B. S. (1994) Conversion of alpha 1-antichymotrypsin into a human neutrophil elastase inhibitor: demonstration of variants with different association rate constants, stoichiometries of inhibition, and complex stabilities. Biochemistry 33, 7627–7633.

(14) Lawrence, D. A., Olson, S. T., Muhammad, S., Day, D. E., Kvassman, J. O., Ginsburg, D., and Shore, J. D. (2000) Partitioning of serpin-proteinase reactions between stable inhibition and substrate cleavage is regulated by the rate of serpin reactive center loop insertion into β-sheet A. J. Biol. Chem. 275, 5839–5844.

(15) Hopkins, P. C. R., Carrell, R. W., and Stone, S. R. (1993) Effects of mutations in the hinge region of serpins. Biochemistry 32, 7650–7657.

(16) Rau, J. C., Beaulieu, L. M., Huntington, J. A., and Church, F. C. (2007) Serpins in thrombosis, hemostasis and fibrinolysis. J. Thromb. Haemost. 5 Suppl 1, 102–15.

(17) Polderdijk, S. G. I., Adams, T. E., Ivanciu, L., Camire, R. M., Baglin, T. P., and Huntington, J. A. (2017) Design and characterization of an APC-specific serpin for the treatment of hemophilia. Blood 129, 105–113.

(18) Owen, M. C., Brennan, S. O., Lewis, J. H., and Carrell, R. W. (1983) Mutation of Antitrypsin to Antithrombin. N. Engl. J. Med. 309, 694–698.

(19) Ehrlich, H. J., Gebbink, R. K., Keijer, J., Linders, M., Preissner, K. T., and Pannekoek, H. (1990) Alteration of serpin specificity by a protein cofactor: Vitronectin endows plasminogen activator inhibitor 1 with thrombin inhibitory properties. J. Biol. Chem. 265, 13029–13035.

(20) Lawrence, D. a, Strandberg, L., Ericson, J., and Ny, T. (1990) Structure-Function Inhibitor Type 1 Studies of the SERPIN Plasminogen Activator. J. Biol. Chem. 265, 20293–20301.

(21) Patston, P. A., Roodi, N., Schifferli, J. A., Bischoff, R., Courtney, M., and Schapira, M. (1990) Reactivity of α1-antitrypsin mutants against proteolytic enzymes of the kallikrein-kinin, complement, and fibrinolytic systems. J. Biol. Chem. 265, 10786–10791.

(22) Hopkins, P. C. R., Crowther, D. C., Carrell, R. W., and Stone, S. R. (1995) Development of a novel recombinant serpin with potential antithrombotic properties. J. Biol. Chem. 270, 11866–11871.

(23) Chaillan-Huntington, C. E., Gettins, P. G. W., Huntington, J. A., and Patston, P. A. (1997) The P6-P2 region of serpins is critical for proteinase inhibition and complex stability. Biochemistry 36, 9562–9570.

(24) Djie, M. Z., Stone, S. R., and Le Bonniec, B. F. (1997) Intrinsic specificity of the reactive site loop of α1-antitrypsin, α1-antichymotrypsin, antithrombin III, and protease nexin I. J. Biol. Chem. 272, 16268–16273.

(25) Plotnick, M. I., Schechter, N. M., Wang, Z. M., Liu, X., and Rubin, H. (1997) Role of the P6-P3’ region of the serpin reactive loop in the formation and breakdown of the inhibitory complex. Biochemistry 36, 14601–14608.

(26) Whisstock, J. C., Silverman, G. A., Bird, P. I., Bottomley, S. P., Kaiserman, D., Luke, C. J., Pak, S. C., Reichhart, J. M., and Huntington, J. A. (2010) Serpins flex their muscle: II. Structural insights into target peptidase recognition, polymerization, and transport functions. J. Biol. Chem. 285, 24307–24312.

(27) Qureshi, T., Goswami, S., McClintock, C. S., Ramsey, M. T., and Peterson, C. B. (2016) Distinct encounter complexes of PAI-1 with plasminogen activators and vitronectin revealed by changes in the conformation and dynamics of the reactive center loop. Protein Sci. 25, 499–510.

(28) Gettins, P. G. W., and Olson, S. T. (2009) Exosite determinants of serpin specificity. J. Biol. Chem. 284, 20441–20445.

(29) Porebski, B. T., Keleher, S., Hollins, J. J., Nickson, A. A., Marijanovic, E. M., Borg, N. A., Costa, M. G. S., Pearce, M. A., Dai, W., Zhu, L., Irving, J. A., Hoke, D. E., Kass, I., Whisstock, J. C., Bottomley, S. P., Webb, G. I., McGowan, S., and Buckle, A. M. (2016) Smoothing a rugged protein folding landscape by sequence-based redesign. Sci. Rep. 6, 33958.

(30) Yang, L., Irving, J. A., Dai, W., Aguilar, M., and Bottomley, S. P. (2018) Probing the folding pathway of a consensus serpin using single tryptophan mutants. Sci. Rep. 1–15.

(31) Zhou, A., Faint, R., Charlton, P., Dafforn, T. R., Carrells, R. W., and Lomas, D. A. (2001) Polymerization of Plasminogen Activator Inhibitor-1. J. Biol. Chem. 276, 9115–9122.

(32) Dafforn, T. R., Mahadeva, R., Elliott, P. R., Sivasothy, P., and Lomas, D. A. (1999) A Kinetic Mechanism for the Polymerization of α 1-Antitrypsin. J. Biol. Chem. 274, 9548–9555.

(33) Belorgey, D., Hägglöf, P., Onda, M., and Lomas, D. A. (2010) pH-dependent stability of neuroserpin is mediated by histidines 119 and 138; Implications for the control of β-sheet a and polymerization. Protein Sci. 19, 220–228.

(34) Horvath, A. J., Lu, B. G. C., Pike, R. N., and Bottomley, S. P. (2011) Methods to measure the kinetics of protease inhibition by serpins. Methods Enzymol. 1st ed. Elsevier Inc.

(35) Buck, M. J., and Atchley, W. R. (2005) Networks of Coevolving Sites in Structural and Functional Domains of Serpin Proteins. Mol. Biol. Evol. 22, 1627–1634.

(36) Irving, J. A., Pike, R. N., Lesk, A. M., and Whisstock, J. C. (2000) Phylogeny of the serpin superfamily: Implications of patterns of amino acid conservation for structure and function. Genome Res. 10, 1845–1864.

(37) Whisstock, J. C., Skinner, R., Carrell, R. W., and Lesk, a M. (2000) Conformational changes in serpins: I. The native and cleaved conformations of alpha(1)-antitrypsin. J. Mol. Biol. 296, 685–699.

(38) Djie, M. Z., Stone, S. R., and Le Bonniec, B. F. (1997) Intrinsic specificity of the reactive site loop of α1-antitrypsin, α1-antichymotrypsin, antithrombin III, and protease nexin I. J. Biol. Chem. 272, 16268–16273.

(39) Wei, A., Rubin, H., Cooperman, B. S., and Christianson, D. W. (1994) Crystal structure of an uncleaved serpin reveals the conformation of an inhibitory reactive loop. Nat. Struct. Mol. Biol. 1, 251–258.

(40) Schreiber, G., and Fersht, A. R. (1996) Rapid, electrostatically assisted association of proteins. Nat. Struct. Biol. 3, 427–431.

(41) Schreiber, G., and Fersht, A. R. (1993) Interaction of Barnase with Its Polypeptide Inhibitor Barstar Studied by Protein Engineering. Biochemistry 32, 5145–5150.

(42) Plotnick, M. I., Rubin, H., and Schechter, N. M. (2002) The Effects of Reactive Site Location on the Inhibitory Properties of the Serpin α 1-Antichymotrypsin. J. Biol. Chem. 277, 29927–29935.

(43) Liu, L., Mushero, N., Hedstrom, L., and Gershenson, A. (2007) Short-lived protease serpin complexes: partial disruption of the rat trypsin active site. Protein Sci. 16, 2403–2411.

(44) Liu, L., Mushero, N., Hedstrom, L., and Gershenson, A. (2006) Conformational distributions of protease-serpin complexes: A partially translocated complex. Biochemistry 45, 10865–10872.

(45) Gong, L., Liu, M., Zeng, T., Shi, X., Yuan, C., Andreasen, P. A., and Huang, M. (2015) Crystal structure of the Michaelis complex between tissue-type plasminogen activator and plasminogen activators inhibitor-1. J. Biol. Chem. 290, 25795–25804.

(46) Lin, Z., Jiang, L., Yuan, C., Jensen, J. K., Zhang, X., Luo, Z., Furie, B. C., Furie, B., Andreasen, P. a, and Huang, M. (2011) Structural basis for recognition of urokinase-type plasminogen activator by plasminogen activator inhibitor-1. J. Biol. Chem. 286, 7027–32.

(47) Madison, E. L., Goldsmith, E. J., Gerard, R. D., Gething, M. J., Sambrook, J. F., and Bassel-Duby, R. S. (1990) Amino acid residues that affect interaction of tissue-type plasminogen activator with plasminogen activator inhibitor 1. Proc. Natl. Acad. Sci. U. S. A. 87, 3530–3.

(48) Madison, E. L., Goldsmith, E. J., Gerard, R. D., Gething, M. J., and Sambrook, J. F. (1989) Serpin-resistant mutants of human tissue-type plasminogen activator. Nature 339, 721–724.

(49) Kass, I., Knaupp, A. S., Bottomley, S. P., and Buckle, A. M. (2012) Conformational properties of the disease-causing Z variant of α1-antitrypsin revealed by theory and experiment. Biophys. J. 102, 2856–2865.

(50) Hughes, V. A., Meklemburg, R., Bottomley, S. P., and Wintrode, P. L. (2014) The Z mutation alters the global structural dynamics of α1-antitrypsin. PLoS One 9, e102617.

(51) Levina, V., Dai, W., Knaupp, A. S., Kaiserman, D., Pearce, M. C., Cabrita, L. D., Bird, P. I., and Bottomley, S. P. (2009) Expression, purification and characterization of recombinant Z α1-Antitrypsin-The most common cause of α1-Antitrypsin deficiency. Protein Expr. Purif. 68, 226–232.

(52) Huang, X., Zheng, Y., Zhang, F., Wei, Z., Wang, Y., Carrell, R. W., Read, R. J., Chen, G. Q., and Zhou, A. (2016) Molecular mechanism of Z α1-antitrypsin deficiency. J. Biol. Chem. 291, 15674–15686.

(53) Lee, K. N., Im, H., Kang, S. W., and Yu, M. H. (1998) Characterization of a human alpha1-antitrypsin variant that is as stable as ovalbumin. J. Biol. Chem. 273, 2509–2516.

(54) Seo, E. J., Lee, C., and Yu, M. H. (2002) Concerted regulation of inhibitory activity of α1-antitrypsin by the native strain distributed throughout the molecule. J. Biol. Chem. 277, 14216–14220.

(55) Cabrita, L. D., Whisstock, J. C., and Bottomley, S. P. (2002) Probing the role of the F-helix in serpin stability through a single tryptophan substitution. Biochemistry 41, 4575–4581.

(56) Cabrita, L. D., Dai, W., and Bottomley, S. P. (2004) Different conformational changes within the F-helix occur during serpin folding, polymerization, and proteinase inhibition. Biochemistry 43, 9834–9839.

(57) Tsutsui, Y., Liu, L., Gershenson, A., and Wintrode, P. L. (2006) The Conformational Dynamics of a Metastable Serpin Studied by Hydrogen Exchange and Mass Spectrometry. Biochemistry 45, 6561–6569.

(58) Cabrita, L. D., Dai, W., and Bottomley, S. P. (2006) A family of E. coli expression vectors for laboratory scale and high throughput soluble protein production. BMC Biotechnol. 6, 12.

(59) Battye, T. G. G., Johnson, O., Powell, H. R., and Leslie, A. G. W. (2011) iMOSFLM: a new graphical interface for diffraction-image processing with MOSFLM research papers. Acta Crystallogr. D Biol. Crystallogr 67, 271–281.

(60) Evans, P. (2006) Scaling and assessment of data quality. Acta Crystallogr. Sect. D Biol. Crystallogr. 62, 72–82.

(61) Winn, M. D., Ballard, C. C., Cowtan, K. D., and Dodson, E. J. (2011) Overview of the CCP 4 suite and current developments. Acta Crystallogr. Sect. D Biol. Crystallogr. 67, 235–242.

(62) McCoy, A. J. (2007) Phaser crystallographic software. J Appl Crystallogr 40, 658–674.

(63) Adams, P. D. (2010) PHENIX: a comprehensive Python-based system for macromolecular structure solution. Acta Crystallogr. Sect. D Biol. Crystallogr. 66, 213–221.

(64) Emsley, P., and Cowtan, K. (2004) COOT: Model-Building Tools for Molecular Graphics. Acta Crystallogr. D Biol. Crystallogr 60, 2126–2132.

(65) Eswar, N., Webb, B., Marti-Renom, M. A., Madhusudhan, M. S., Eramian, D., Shen, M. M., Pieper, U., Sali, A., and Marti-Renom, M. A. (2002) Comparative protein structure modeling using Modeller, in Current Protocols in Bioinformatics, pp 5–6.

(66) Patschull, A. O. M., Segu, L., Nyon, M. P., Lomas, D. A., Nobeli, I., Barrett, T. E., and Gooptu, B. (2011) Therapeutic target-site variability in α 1-antitrypsin characterized at high resolution. Acta Crystallogr. Sect. F Struct. Biol. Cryst. Commun. 67, 1492–1497.

(67) Schrodinger, L. The Pymol Molecular Graphics System, Version 2.0.4.

(68) Humphrey, W., Dalke, A., and Schulten, K. (1996) VMD: Visual molecular dynamics. J. Mol. Graph. 14, 33–38.

(69) Mcgibbon, R. T., Beauchamp, K. A., Schwantes, C. R., Wang, L., Hern, C. X., Herrigan, M. P., Lane, T. J., Swails, J. M., and Pande, V. S. (2014) MDTraj: a modern, open library for the analysis of molecular dynamics trajectories MDTraj: a modern, open library for the analysis of molecular dynamics trajectories. Biorxiv.Org 109, 0–2.

(70) Baker, N. A., Sept, D., Joseph, S., Holst, M. J., and McCammon, J. A. (2001) Electrostatics of nanosystems: Application to microtubules and the ribosome. Proc. Natl. Acad. Sci. 98, 10037–10041.

(71) Jurrus, E., Engel, D., Star, K., Monson, K., Brandi, J., Felberg, L. E., Brookes, D. H., Wilson, L., Chen, J., Liles, K., Chun, M., Li, P., Gohara, D. W., Dolinsky, T., Konecny, R., Koes, D. R., Nielsen, J. E., Head-Gordon, T., Geng, W., Krasny, R., Wei, G.-W., Holst, M. J., McCammon, J. A., and Baker, N. A. (2018) Improvements to the APBS biomolecular solvation software suite. Protein Sci. 27, 112–128.

(72) Ye, S., Cech, a L., Belmares, R., Bergstrom, R. C., Tong, Y., Corey, D. R., Kanost, M. R., and Goldsmith, E. J. (2001) The structure of a Michaelis serpin-protease complex. Nat. Struct. Biol. 8, 979–983.

(73) Dolinsky, T. J., Nielsen, J. E., McCammon, J. A., and Baker, N. A. (2004) PDB2PQR: An automated pipeline for the setup of Poisson-Boltzmann electrostatics calculations. Nucleic Acids Res. 32, W665–W667.

(74) Søndergaard, C. R., Olsson, M. H. M., and Rostkowski Michałand Jensen, J. H. (2011) Improved Treatment of Ligands and Coupling Effects in Empirical Calculation and Rationalization of pKa Values. J Chem Theory Comput 7, 2284–2295.

(75) Jorgensen, W. L., Chandrasekhar, J., Madura, J. D., Impey, R. W., Klein, M. L., Jorgensen, W. L., Chandrasekhar, J., Madura, J. D., Impey, R. W., and Klein, M. L. (2001) Comparison of simple potential functions for simulating liquid water Comparison of simple potential functions for simulating liquid water. J. Chem. Phys. 926, 926–935.

(76) Suk Joung, I., and Cheatham, T. E. (2008) Determination of Alkali and Halide Monovalent Ion Parameters for Use in Explicitly Solvated Biomolecular Simulations. J. Phys. Chem. B 112, 9020–9041.

(77) Maier, J. A., Martinez, C., Kasavajhala, K., Wickstrom, L., Hauser, K. E., and Simmerling, C. (2015) ff14SB: Improving the Accuracy of Protein Side Chain and Backbone Parameters from ff99SB. J. Chem. Theory Comput. 11, 3696–3713.

(78) Li, P., Roberts, B. P., Chakravorty, D. K., and Merz, K. M. (2013) Rational design of particle mesh ewald compatible lennard-jones parameters for +2 metal cations in explicit solvent. J. Chem. Theory Comput. 9, 2733–2748.

(79) Case DA, Berryman JT, Betz RM, Cai Q, Cerutti DS, Cheatham TE, et. al. (2014) Amber 14.

(80) Edgar, R. C. (2004) MUSCLE: Multiple sequence alignment with high accuracy and high throughput. Nucleic Acids Res. 32, 1792–1797.

(81) Katoh, K., and Standley, D. M. (2013) MAFFT multiple sequence alignment software version 7: Improvements in performance and usability. Mol. Biol. Evol. 30, 772–780.

(82) Landau, M., Mayrose, I., Rosenberg, Y., Glaser, F., Martz, E., Pupko, T., and Ben-Tal, N. (2005) ConSurf 2005: The projection of evolutionary conservation scores of residues on protein structures. Nucleic Acids Res. 33, 299–302.

(83) Ashkenazy, H., Abadi, S., Martz, E., Chay, O., Mayrose, I., Pupko, T., and Ben-Tal, N. (2016) ConSurf 2016: an improved methodology to estimate and visualize evolutionary conservation in macromolecules. Nucleic Acids Res. 44, W344–W350.

(84) Kass, I., and Horovitz, A. (2002) Mapping pathways of allosteric communication in GroEL by analysis of correlated mutations. Proteins Struct. Funct. Genet. 48, 611–617.

(85) Lockless, S. W., and Ranganathan, R. (1999) Evolutionarily conserved pathways of energetic connectivity in protein families. Science 286, 295–9.

(86) Dekker, J. P., Fodor, A., Aldrich, R. W., and Gary, Y. (2004) A perturbation-based method for calculating explicit likelihood of evolutionary co-variance in multiple sequence alignments. Bioinformatics 20, 1565–1572.

(87) McGibbon, R. T., Beauchamp, K. A., Harrigan, M. P., Klein, C., Swails, J. M., Hernández, C. X., Schwantes, C. R., Wang, L.-P., Lane, T. J., and Pande, V. S. (2015) MDTraj: A Modern Open Library for the Analysis of Molecular Dynamics Trajectories. Biophys. J. 109, 1528–32.

(88) Hunter, J. D. (2007) Matplotlib: A 2D graphics environment. Comput. Sci. Eng. 9, 99–104.

(89) Campello, R. J. G. B., Moulavi, D., and Sander, J. (2013) Density-Based Clustering Based on Hierarchical Density Estimates, in Antimicrobial agents and chemotherapy, pp 160–172.

(90) McInnes, L., Healy, J., and Astels, S. (2017) hdbscan: Hierarchical density based clustering. J. Open Source Softw. 2, 11–12.

(91) Parra, R. G., Schafer, N. P., Radusky, L. G., Tsai, M. Y., Guzovsky, A. B., Wolynes, P. G., and Ferreiro, D. U. (2016) Protein Frustratometer 2: a tool to localize energetic frustration in protein molecules, now with electrostatics. Nucleic Acids Res. 44, W356–W360.

